# Three quantitative trait loci explain more than 60% of phenotypic variation for chill coma recovery time in *Drosophila ananassae*

**DOI:** 10.1101/676262

**Authors:** Annabella Königer, Saad Arif, Sonja Grath

## Abstract

Ectothermic species such as insects are particularly vulnerable to climatic fluctuations. Nevertheless, many insects that evolved and diversified in the tropics have successfully colonized temperate regions all over the globe. To shed light on the genetic basis of cold tolerance in such species, we conducted a quantitative trait locus (QTL) mapping experiment for chill coma recovery time (CCRT) in *Drosophila ananassae*, a cosmopolitan species that has expanded its range from tropical to temperate regions.

We created a mapping population of recombinant inbred advanced intercross lines (RIAILs) from two founder strains with diverging CCRT phenotypes. The RIAILs were phenotyped for their CCRT and, together with the founder strains, genotyped for polymorphic markers with double-digest restriction site-associated DNA (ddRAD) sequencing. Using a hierarchical mapping approach that combined standard interval mapping and a multiple-QTL model, we mapped three QTL which altogether explained 64% of the phenotypic variance. For two of the identified QTL, we found evidence of epistasis. To narrow down the list of cold tolerance candidate genes, we cross-referenced the QTL intervals with genes that we previously identified as differentially expressed in response to cold in *D. ananassae*, and with thermotolerance candidate genes of *D. melanogaster*. Among the 58 differentially expressed genes that were contained within the QTL, *GF15058* showed a significant interaction of the CCRT phenotype and gene expression. Further, we identified the orthologs of four *D. melanogaster* thermotolerance candidate genes, *MtnA*, *klarsicht*, *CG5246* (*D.ana*/*GF17132*) and *CG10383* (*D.ana*/*GF14829*) as candidates for cold tolerance in *D. ananassae*.

## Introduction

Temperature is one of the major factors that influence the geographical distribution and abundance of ectothermic species. Physiological mechanisms to regulate body temperature are usually limited in ectotherms and resilience towards temperature extremes often determines the species fate upon climate change or range expansion. *Drosophila* spp. have successfully mastered such thermal challenges as they colonized temperate regions all over the globe and are now present on all of the earth’s continents except Antarctica (Lachaise et al., 1988). By far the most prominent example of the genus is *Drosophila melanogaster*, which originated in sub-Saharan Africa, colonized temperate regions after the last glaciation about 15,000 years ago and nowadays has a worldwide distribution (David and Capy, 1988; Stephan and Li, 2007). Cold tolerance in this species has a highly polygenic basis (von Heckel et al., 2016; MacMillan et al., 2016) and adaptation to local temperatures required simultaneous selection at multiple loci (Morgan and Mackay, 2006; Svetec et al., 2011).

Previously, we examined the cold tolerance of *Drosophila ananassae* (Königer and Grath, 2018), a tropical species which originated in South-East-Asia (Das et al., 2004). During the past 18.000 years, *D. ananassae* expanded from its ancestral range to temperate regions and has nowadays a quasi-cosmopolitan distribution (Das et al., 2004; Tobari, 1993) We measured cold tolerance by means of a test for chill coma recovery time (CCRT), which is defined as the time the flies need to stand on their legs after a cold-induced coma (David et al., 1998). There was substantial variation in CCRT among fly strains that were derived from a population of the ancestral species range in Bangkok, Thailand (Königer and Grath, 2018). Most strikingly, the difference in the phenotype within this single population was large if compared to within-population variance in *D. melanogaster* (von Heckel et al., 2016). However, in *D. ananassae*, only two genes, *GF15058* and *GF14647*, reacted to the cold shock in a phenotype-specific, i.e., they showed a significant interaction of phenotype and genotype.

Here, we report the results of a genome-wide scan for quantitative trait loci (QTL) influencing CCRT in *D. ananassae.* To gain better insight into the genetic architecture of cold tolerance in this species, we generated a mapping population of recombinant inbred advanced intercross lines (RIAILs) from the most cold-tolerant strain and the most cold-sensitive strain of the Bangkok population. By combining double-digest restriction site-associated DNA sequencing (ddRAD) markers and a hierarchical mapping approach, we identified three QTL of large effect which altogether explain 64% of the variance in the phenotype. We further combined the present results with lists of genes that are differentially expressed in response to the cold shock in *D. ananassae* (Königer and Grath, 2018) and *D. melanogaster* (von Heckel et al., 2016).

Both species belong to the Melanogaster group and shared a common ancestor around 15-20 million years ago (Drosophila 12 Genomes Consortium et al., 2007). *D. melanogaster* expanded its range from Sub-Saharan Africa to temperate regions in Europe about 16,000 years ago (Stephan and Li, 2007). Our approach allowed us to narrow down the list of potentially causal genes for cold tolerance and to uncover common evolutionary patterns among species from completely independent phylogenetic lineages that have expanded their thermal ranges and became successful human commensals.

## Materials and Methods

### Mapping population

All flies used in this study were raised on standard cornmeal molasses medium, at constant room temperature (22 ± 1°C) and at a 14:10 h light:dark cycle (details of the food recipe can be found in (Königer and Grath, 2018)). The two fly strains (Fast and Slow) that were used as founders for the mapping population were collected in 2002 in Bangkok, Thailand, and established as isofemale strains (Das et al., 2004). Recombinant Inbred Advanced Intercross Lines (RIAILs) were generated as follows (Figure 1): two initial crosses between the two parental strains were set up (Fast males × Slow females and Slow males × Fast females). Individuals from both F1 generations were mixed and allowed to mate freely with each other. Up to generation F4, intercrossing was continued in the form of mass breedings. In generation F4, 360 mating pairs were set up in separate vials to allow for one more generation of intercrossing and to initiate the inbred strains. From generation F5, full-sibling inbreeding was carried out by mating brother-sister pairs for five subsequent generations. Throughout all generations (P – F10), the parents were removed before the offspring hatched to avoid back-crosses. From generation F10 on, RIAILs were kept at low density in 50 ml vials.

**Figure 1.**
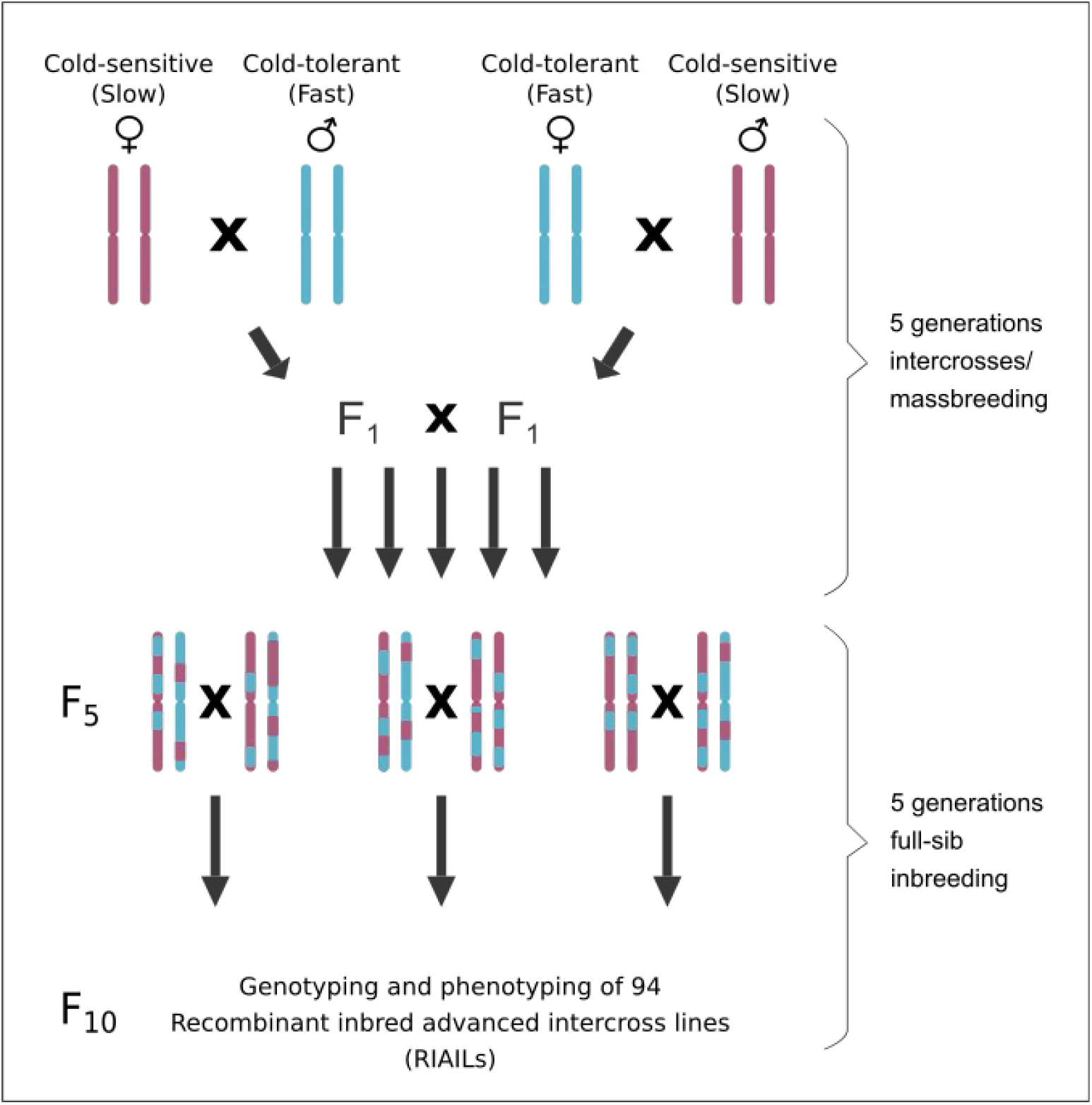
Crossing scheme for the generation of the RIAIL mapping population. Drawings of single chromosome pairs were used as representatives for the full genome. An initial, reciprocal cross between the cold-sensitive founder strain Slow (shown in red) and the cold-tolerant founder strain Fast (shown in blue) was set up to generate the heterozygous F1 generation. Intercrosses were continued in the form of massbreedings until generation F4, where single mating pairs were picked to allow for one more generation of intercrossing and to initiate inbreeding. From generation F5, full-sibling inbreeding was carried out for five subsequent generations. Throughout all generations (P – F10), the parents were removed before the offspring hatched to avoid back-crosses.

### Test for chill coma recovery time (CCRT)

CCRT was measured for flies of 4 – 6 days of age as described previously (Königer and Grath, 2018). For the two founder strains Fast and Slow, CCRT was measured for males and females separately. For the RIAILs, only female flies were phenotyped. All female flies were collected and phenotyped as virgins. In brief, collection and sex-separation were carried out under light CO_2_-anesthesia, whereby ten flies from the same sex and strain were collected into a 50 ml vial containing 10 ml of cornmeal molasses medium. At the age of 4–6 days, the flies were transferred without anesthesia into new vials without food. For the cold shock, the vials were placed in an ice water bath (0 ± 0.5 °C) for exactly 3 h. Back at room temperature (22 ± 1 °C), CCRT was monitored in 2 min intervals for the duration of 90 min. Flies that were still not standing after 90 min were assigned a recovery time of 92 min. Flies that died during the experiment (< 1 %) were excluded from the analysis. On average, we tested 40 female individuals per RIAIL and 100 individuals per founder strain and sex.

### DNA extraction and sequencing

DNA was extracted from 94 RIAILs and the two parental strains with the DNeasy® Blood & Tissue Kit (QIAGEN, Hilden, Germany). For each strain, 10 virgin female individuals were pooled. DNA concentration and purity were assessed with a spectrophotometer (NanoDrop® ND 1000, VWR International, Radnor, PA, USA). Library preparation and double-digest restriction site-associated DNA sequencing (ddRAD-seq) was carried out by an external sequencing service (ecogenics GmbH, Balgach, Switzerland) in the following way: DNA was double-digested with EcoRI and MseI and ligated to respective adapters comprising EcoRI and MseI restriction overhangs. Molecular identifier tags were added by polymerase chain reaction. The individual sample libraries were pooled, and the resulting library pools were size-selected for fragments between 500-600 bp with gel electrophoresis and extraction of the respective size range. The resulting size selected library pools were sequenced on a NextSeqTM 500 Sequencing System (Illumina, San Diego, CA), producing single-ended reads of 75bp length. Demultiplexing and trimming from Illumina adapter residuals was also carried out by the external service.

### Marker catalog construction and data curation

The software pipeline Stacks (version 1.45) (Catchen et al., 2011) was used to analyze the sequence data and to identify markers. First, to examine the quality of the sequence reads, the *process_radtags* program was run in Stacks, applying a sliding window size of 50% of the read length (*-w 0.5*) to filter out reads which drop below a 99% probability of being correct (Phred score < 20) (*-s 20*). Second, the processed reads of each sample were mapped to the *D. ananassae* reference genome (FlyBase release 1.05 (Attrill et al., 2016) with NextGenMap (version 0.5.0) (Sedlazeck et al., 2013). Third, the mapped reads were converted to bam format, sorted and indexed with samtools (version 0.1.18) (Li et al., 2009). Fourth, the *ref_map.pl* wrapper program was run in Stacks, which executes the Stacks core pipeline by running each of the Stacks components individually. In brief, *pstacks* assembled RAD loci for each sample, *cstacks* created a catalog of RAD loci from the two parental samples to create a set of all possible alleles expected in the mapping population and *sstacks* matched all RIAIL samples against the catalog. The *genotypes* program was executed last, applying automated corrections to the data (*-c*) to correct for false-negative heterozygote alleles. Only those loci which were present in at least 80% of the samples were exported (*-r 75*). Fifth, we applied additional corrections to the catalog by running the *rxstacks* program with the following filtering settings: non-biological haplotypes unlikely to occur in the population were pruned out (*--prune_haplo*), SNPs were recalled once sequencing errors were removed using the bounded SNP model (*--model_type_bounded*) with an error rate of 10% *(--bound_high 0.1*), and catalog loci with an average log likelihood less than –200 were removed (*--lnl_lim −200.00*). Sixth, *cstacks* and *sstacks* and *genotypes* (*-r 75*) were rerun to rebuild, match and export a new catalog with the filtered SNP calls. *Load_radtags.pl* and *index_radtags.pl* were used to upload and index the new catalog to a MySQL database. Seventh, a custom R script was used to remove markers with extreme values of residual heterozygosity within RIAILs, using cutoffs based on our inbreeding scheme (> 15% and < 35%) (Falconer and Mackay, 1996) and to remove markers with an allele frequency drift < 10% from further analysis. Eighth, the MySQL database was used to manually check the markers for errors. A total of 1,400 markers were included in the downstream analysis.

### Genetic map construction

Genetic map construction was conducted with R/qtl (version 1.42) (Broman et al., 2003). The function *countXO* was used to remove seven RIAILs with > 200 crossover events. One more RIAIL was removed due to a low number of genotyped markers (< 700). The downstream analysis included 1,400 markers and 86 RIAIL-samples (Supplementary file 1: Table S3). Markers were partitioned into linkage groups based on a logarithm of the odds (LOD) score threshold of 8 and a maximum recombination frequency (rf) of 0.35, assuming a sequencing error rate of 1%. Map distances were calculated using the Haldane map function. As a sanity check, the functions *plotRF* and *checkAlleles* were used to test for potentially switched alleles and linkage groups were visually validated (based on rf and LOD scores).

### Analysis of quantitative trait loci (QTL)

QTL mapping was conducted with R/qtl (version 1.42) (Broman et al., 2003). Prior to mapping, the genotype probabilities between marker positions were calculated with the function *calc.genoprob* on a maximum grid size of 1 cM. To identify major QTL, standard interval mapping was performed using the Expectation Maximization (EM) algorithm as implemented with the *scanone* function. The results are expressed as a LOD score (Sen and Churchill, 2001). Significance thresholds were calculated with 1,000 genome-wide permutations. The initial single-QTL scan was extended with a more complex, two-dimensional scan using Haley-Knott-Regression as implemented with the *scantwo* function. Significance thresholds were again calculated with 1,000 genome-wide permutations.

To screen for additional QTL, estimate QTL effects and refine QTL positions, multiple-QTL mapping (MQM) was performed (Arends et al., 2010). Here, missing genotypes were simulated from the joint distribution using a Hidden Markov model with 1,000 simulation replicates and an assumed error rate of 1% as implemented with the *sim.geno* function. The MQM model was identified with a forward selection/backward elimination search algorithm as implemented with the *stepwise* function, with the model choice criterion being penalized LOD scores. The penalties were derived on the basis of the significance permutations from the two-dimensional genome scan. To estimate the support interval for each identified QTL, an approximate 95% Bayesian credible interval was calculated as implemented by the *bayesint* function. Gene annotations for QTL intervals were downloaded from FlyBase (Attrill et al., 2016) and screened for enriched GO terms and KEGG pathways with DAVID (version 6.8) (Huang et al., 2009). Enrichment was calculated against the background of all annotated genes (Attrill et al., 2016) using default settings (EASE-score of 0.1 after multiple testing correction according to Benjamini-Hochberg (Benjamini and Hochberg, 1995)). In addition, we cross-referenced the QTL gene lists with lists of differentially expressed genes from a previously conducted transcriptome analysis, where we compared gene expression among cold-tolerant and cold-sensitive fly strains from the Bangkok population (including the two parental founder strains used in this study) in response to the 3 h cold shock at 0°C (Königer and Grath, 2018). Moreover, the transcriptome analysis also comprises lists of differentially expressed genes of cold-tolerant and cold-sensitive fly strains of *Drosophila melanogaster* in response to a cold shock (von Heckel et al., 2016), allowing us to compare expression regulation of orthologous genes within the QTL regions among these two *Drosophila* species.

### Data Availability

The sequence data have been deposited in NCBI’s Sequence Read Archive and are accessible through series accession number PRJNA544044. Supplementary material is deposited at figshare.

## Results

### Chill coma recovery time (CCRT) Phenotype

The average CCRT of the cold-tolerant founder strain (Fast) was 29.29 min for females and 30.13 min for males. CCRT of the cold-sensitive founder strain (Slow) was 63.70 min for females and 53.92 min for males (Figure 2, Supplementary file 1: Table S1). The difference in CCRT between the Fast strain and the Slow strain was significant for males (Welch’s t-test, *P*-value < 2.2×10^−16^) and females (Welch’s t-test, *P*-value < 2.2×10^−16^). The average CCRT of the RIAILs ranged from 27.60 min to 83.03 min (Figure 3, Supplementary file 1: Table S2).

**Figure 2.**
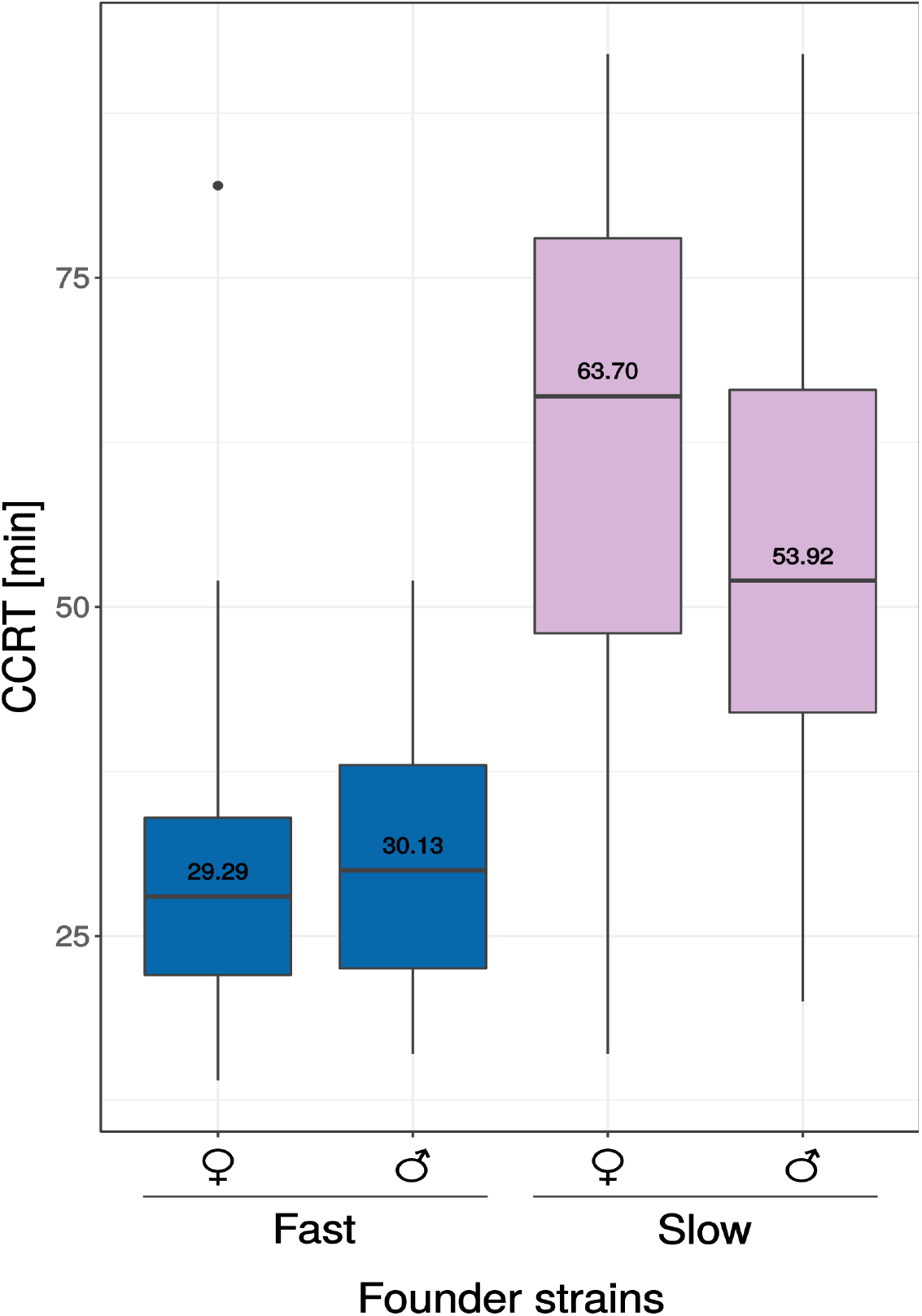
Chill coma recovery time (CCRT) of 4-6 day old flies of two strains of *D. ananassae* from Bangkok (Thailand) that were used as founder strains for the mapping population. In both sexes, the Fast strain recovered significantly faster than the Slow strain (Welch’s t-test, *P*-value < 2.2×10^−16^).

**Figure 3.**
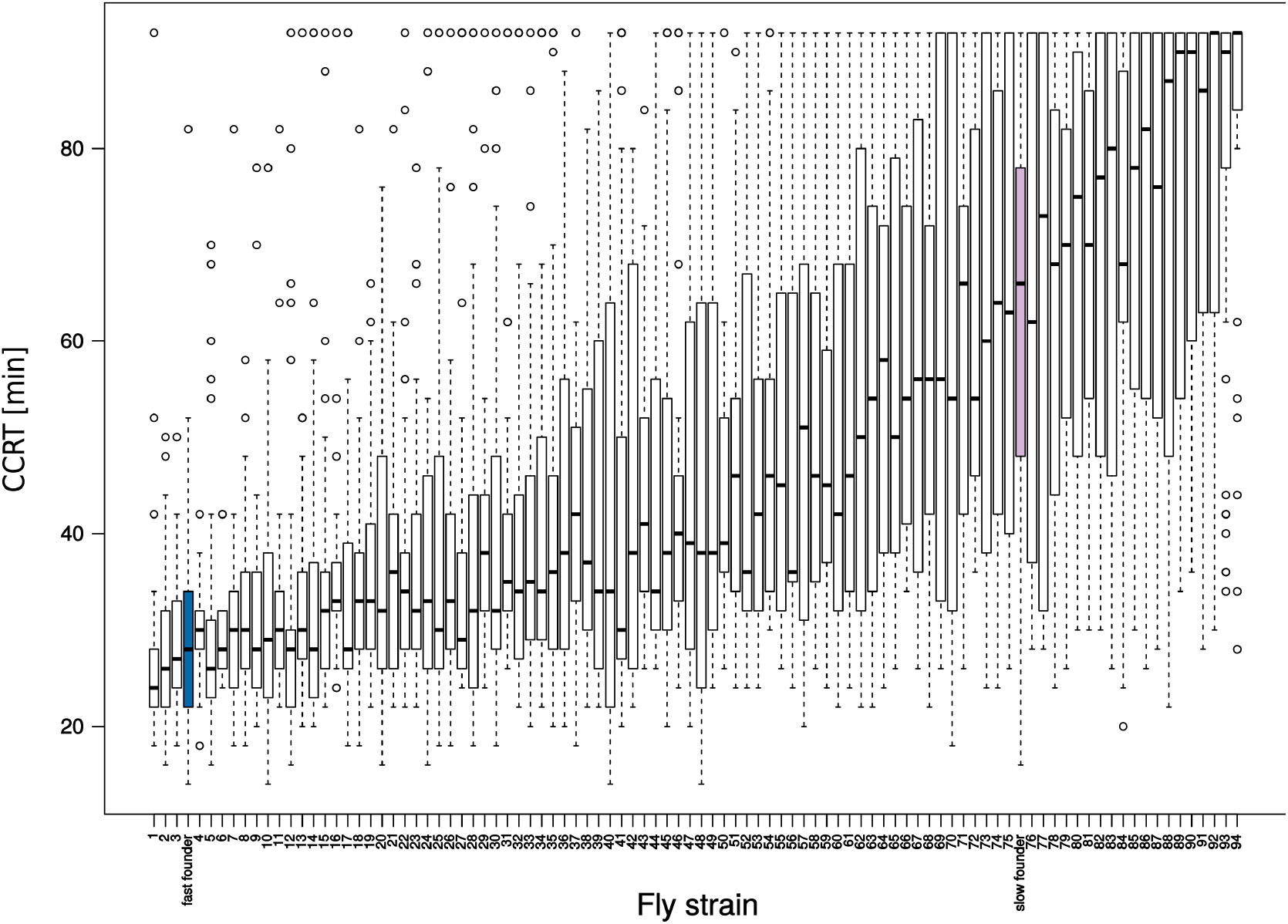
Chill coma recovery time (CCRT) of 4 – 6 day old virgin female flies of the 94 recombinant inbred advanced intercross lines (RIAILs) is displayed with white bars. CCRT of the two founder strains is displayed in blue (Fast founder) and pink (Slow founder) (see also Figure 2). The RIAILs were numbered in ascending order according to their average CCRT.

### Sequencing and genetic map

In total, we obtained 331,867,133 sequence reads with an average of 3,281,450 reads per sample. 0.6% of the total reads (2,074,057) failed the Stacks *process_radtags* quality check and were excluded from the analysis. In each of the samples, > 94% of all reads mapped to the *D. ananassae* reference genome. The Stacks core pipeline matched 5,468 markers to the initial catalog. After additional corrections with the *rxstacks* program, 3,092 markers remained. 1,692 more markers were excluded from this new catalog due to extreme values of heterozygosity and allele frequency drift. Thus, after all filtering steps, a total of 1,400 markers and 86 RIAILs were used for genetic map construction. The markers were partitioned into eight linkage groups (Supplementary file 1: Table S4). The total map length was 962.0 cM, with an average marker spacing of 0.7 cM and a maximum marker spacing of 55.5 cM (Figure 5). Across all samples, 91.6% of the genotypes were available of which 37.4% were homozygous for the cold-tolerant (Fast) allele (FF), 27.9% were heterozygous (FS) and 34.7% were homozygous for the cold-sensitive (Slow) allele (SS).

### One- and two-dimensional genome scans

Interval mapping in the context of a single-QTL model revealed two major areas with LOD peaks which exceeded the permuted 5% significance level (LOD 3.53), one on scaffold 13337 (QTL1) and one on scaffold 13340 (QTL2) (Figure 4). The highest peak on scaffold 13337 was at 6.08 cM (LOD 5.80) and the highest peak on scaffold 13340 was at 80.05 cM (LOD 4.08).

**Figure 4.**
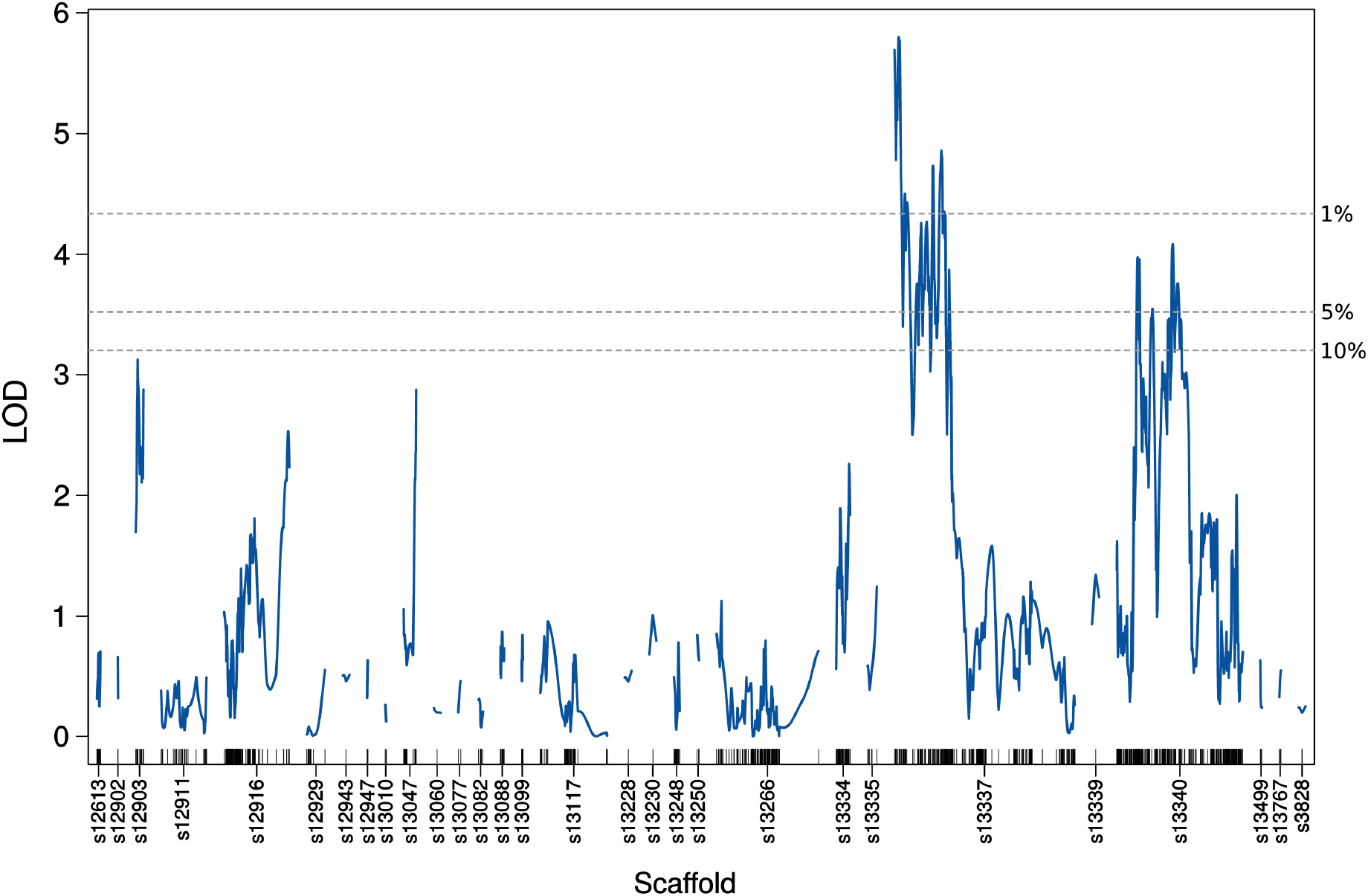
LOD-curves obtained with standard interval mapping reveal two significant QTL, QTL1 on scaffold 13337 and QTL2 on scaffold 14440. Significance thresholds (dotted lines) were calculated with 1,000 genome-wide permutations. The short vertical lines on the X-axis correspond to the marker positions.

The next step was to extend the initial, single-QTL scan with a two-dimensional scan, where we compared two possible models: the full (epistatic) model (H_f1_) which allowed for the possibility of a second QTL and interactions among QTL was compared to the additive model (H_a1_) which allowed for the possibility of a second QTL without interaction. Both the full and the additive model reached maximum LOD scores at the same positions, 7.08 cM on scaffold 13337 and 30.1 cM on scaffold 13340 (Table 1). In comparison to the single-QTL model, we found supporting evidence for the presence of a second QTL under the additive model (lodd.av1 *P*-value = 0.006), but not under the full model (lod.fv1 *P*-value = 0.668). There is no evidence for interaction among the two loci (lod.int *P*-value = 1).

**Table 1.** Results of the two-dimensional genome scan

| **Two-QTL scan** |  |  |  |  |  |  |  |  |
| --- | --- | --- | --- | --- | --- | --- | --- | --- |
| | pos1f<sup>1</sup> | pos2f<sup>1</sup> | lod.full<sup>1</sup> | $P$-value<sup>1*</sup> | lod.fv1<sup>2</sup> | $P$-value<sup>2*</sup> | | |
| s13337:s13340 | 7.08 | 30.1 | 12.6 | 0 | 5.77 | 0.668 |  |  |
| | pos1a<sup>3</sup> | pos2a<sup>3</sup> | lod.add<sup>3</sup> | $P$-value<sup>3*</sup> | lod.av1<sup>4</sup> | $P$-value<sup>4*</sup> | lod.int<sup>5</sup> | $P$-value<sup>5*</sup> |
| s13337:s13340 | 7.08 | 30.1 | 11.4 | 0 | 4.54 | 0.006 | 1.24 | 1 |

1) QTL positions, LOD score and $P$ -value for the full (epistatic) model versus the Null-model
2) LOD score and $P$ -value for the full (epistatic) model versus the Single-QTL-model
3) QTL positions, LOD score and $P$ -value for the additive model versus the Null-model
4) LOD score and $P$ -value for the additive model versus the Single-QTL-model
5) LOD-score and $P$ -value of (full model – additive model) = evidence for interaction
\*) $P$ -values represent the proportion of permutation replicates with LOD scores ≥ the observed

### Multilpe-QTL model

In order to identify possible additional QTL of moderate effect, refine QTL positions, separate linked loci and to estimate QTL effects, we applied a forward selection/backward elimination algorithm with penalized LOD scores and identified a model with three main terms and one interaction term. The overall fit of the model had a LOD score of 19.26 and explained 64.34% of the phenotypic variance (Figure S2, Supplementary file 1: Table S5). In comparison to the one- and two-dimensional genome-scans, there was an additional locus on scaffold 12916 at position 16.7 cM (QTL3) which interacted with one of the previously identified loci, on scaffold 13340 (QTL2) (Table 2, Figures 5, 6 and Figure S3).

**Figure 5.**
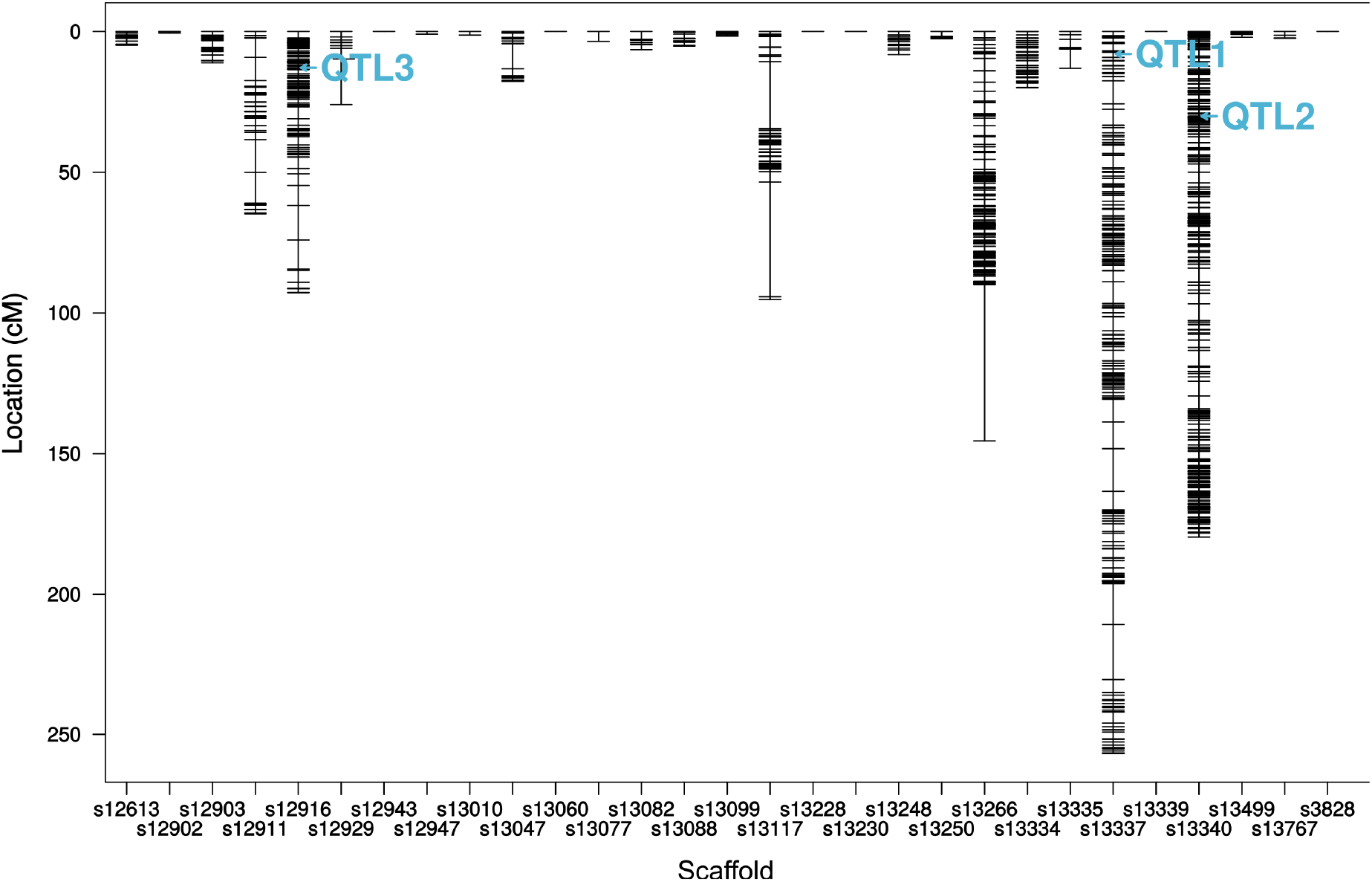
Genetic map with QTL positions as refined with the multiple-QTL model. X-axis = genomic scaffolds. Y-axis = genetic distances in centiMorgan (cM) for markers (short horizontal lines).

**Table 2.** QTL confidence intervals and estimated effects

|  | Scaffold | cytologic position<br>[cM] | cytologic position<br>[bp] | confidence interval<br>[bp] | % variance | additive<br>effect | dominance<br>deviation | genes* |
| --- | --- | --- | --- | --- | --- | --- | --- | --- |
| QTL1 | 13337 | 0.083871 | 0.083871 – 9.233870 | 83.871 – 226.785 | 26.59 | 9.2082 | -2.1719 | 11 |
| QTL2 | 13340 | 30.053110 | 27.51427 – 36.52024 | 5.542.036 – 6.544.039 | 30.44 | 1.7317 | -5.2343 | 138 |
| QTL3 | 12916 | 16.747634 | 7.103214 – 92.933043 | 1.514.827 - 2.696.582 | 19.89 | -0.6307 | -0.8054 | 110 |

QTL positions and effects on the phenotype as estimated with the multiple-QTL model. Confidence intervals were calculated as 95% Bayesian credible intervals.
\* Numbers of protein-coding genes within QTL intervals. Numbers and identifiers for non-coding genes and RNAs are shown in Supplementary file 2.

**Figure 6.**
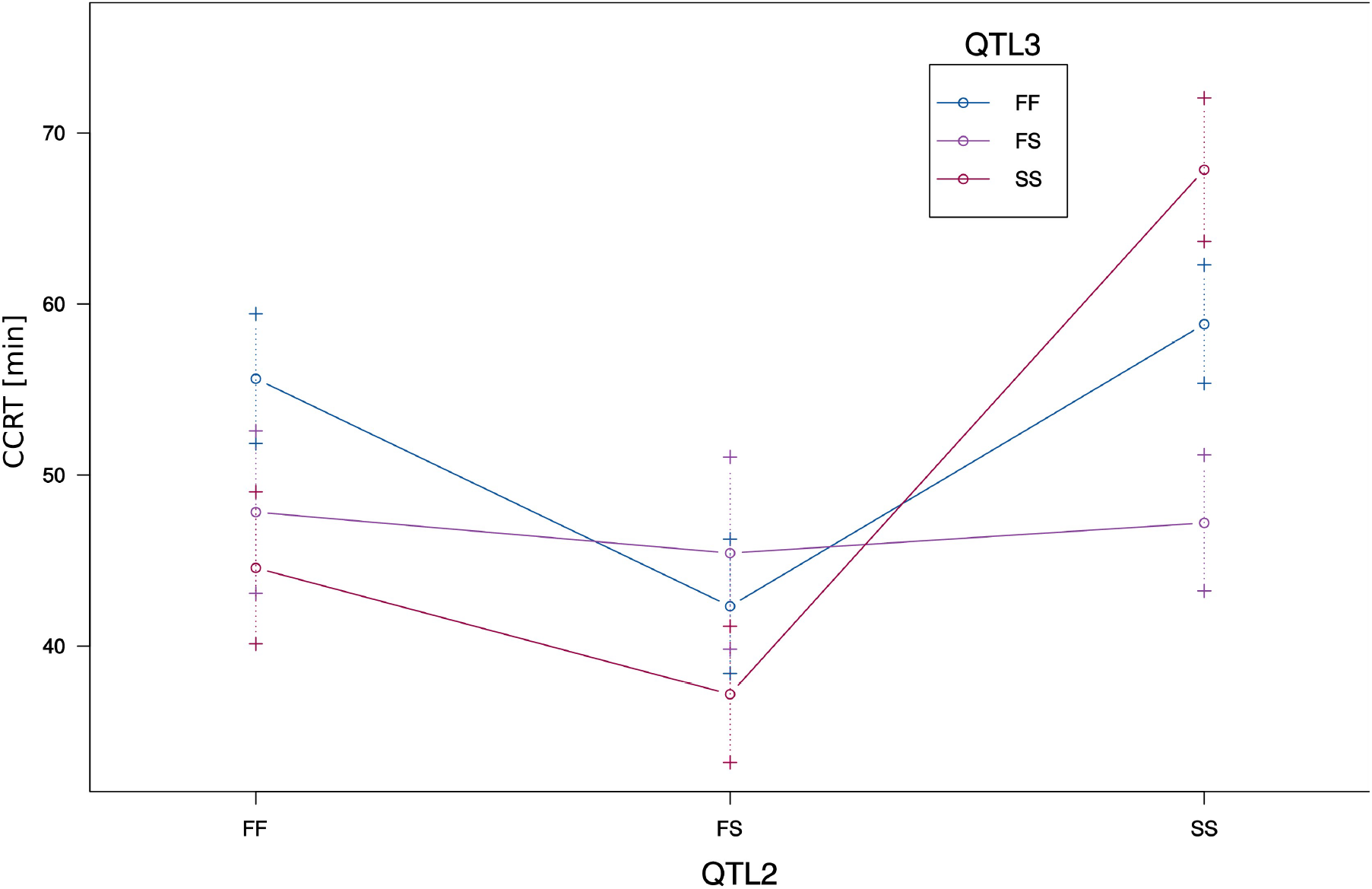
Interaction of QTL2 on scaffold 13340 and QTL3 on scaffold 12916. X-axis = genotypes for QTL2. The genotypes for QTL3 are represented by lines in different colors. Error bars are plotted at +/− 1 SE. F = cold-tolerant parental allele (fast CCR), S = cold-sensitive parental allele (slow CCR).

QTL effects were estimated for additivity ((SS-FF)/2) and deviation from dominance ((FS-(FF+SS)/2), where F denotes the cold-tolerant Fast allele and S denotes the cold-sensitive Slow allele (Table 2, Supplementary file 1: Table S6). QTL3 on scaffold 12916 was a transgressive QTL as the cold-tolerant allele was associated with having a more cold-sensitive phenotype (longer CCRT), resulting in a negative effect size (Figure S1C). For QTL1, the estimated additive effect was positive while the estimated dominance effect was negative. RIAILs homozygous for the cold-tolerant allele had the most cold-tolerant phenotype, RIAILs homozygous for the cold-sensitive allele had the least cold-tolerant phenotype and heterozygote RIAILs had an intermediate phenotype (Figure S1A). The effect estimates for QTL2 went in the same direction as for QTL1. Here, however, the heterozygous phenotype was associated with the most cold-tolerant phenotype (Figure S1B). The more complex relationships of additive and dominance effects for the interaction of QTL3 and QTL2 can be understood best by plotting the interaction of the phenotype and the genotype at both marker positions (Figure 6).

As revealed by the interaction plot (Figure 6), RIAILs homozygous for the cold-sensitive (S) allele at both QTL also had the most cold-sensitive phenotype. The most cold-tolerant phenotype, was reached by those RIAILs which were homozygous for the cold-tolerant allele at QTL2 but homozygous for the cold-sensitive allele at (the transgressive) QTL3 Interestingly, cold tolerance of RIAILs which were heterozygous at QTL3 seemed to be independent from their genotype at QTL2.

The results of a drop-one-term at a time ANOVA indicated strong evidence for all three loci and the interaction of QTL2 and QTL3: for each QTL, the model with the QTL of interest at that particular position was compared to the model with the QTL of interest omitted, while all other QTL positions were fixed at their maximum likelihood estimates (Table 3, Figure S3).

**Table 3.** Summary table for the drop one term ANOVA

| QTL | cytologic position | df | Type III SS | LOD | %Var | F value | *P* (Chi2) | *P* (F) |
| --- | --- | --- | --- | --- | --- | --- | --- | --- |
| 1 | s13337-0.1 | 2 | 4903 | 10.404 | 26.59 | 27.962 | 0 | 8.45E-10 |
| 2 | s13340-30.1 | 6 | 5613 | 11.526 | 30.44 | 10.671 | 0 | 1.52E-08 |
| 3 | s12916-16.7 | 6 | 3667 | 8.277 | 19.89 | 6.972 | 0 | 6.47E-06 |
| 2:3 | s13340-30.1:s12916-16.7 | 4 | 2790 | 6.606 | 15.13 | 7.957 | 0 | 2.11E-05 |

### Candidate gene meta analysis

All three QTL together contained 259 protein-coding genes (Table 4, Supplementary file 2: Tables S1, S2, S5). Among them were 58 genes that we had identified previously as differentially expressed in response to the cold shock (Königer and Grath, 2018, Table 4).

**Table 4.**
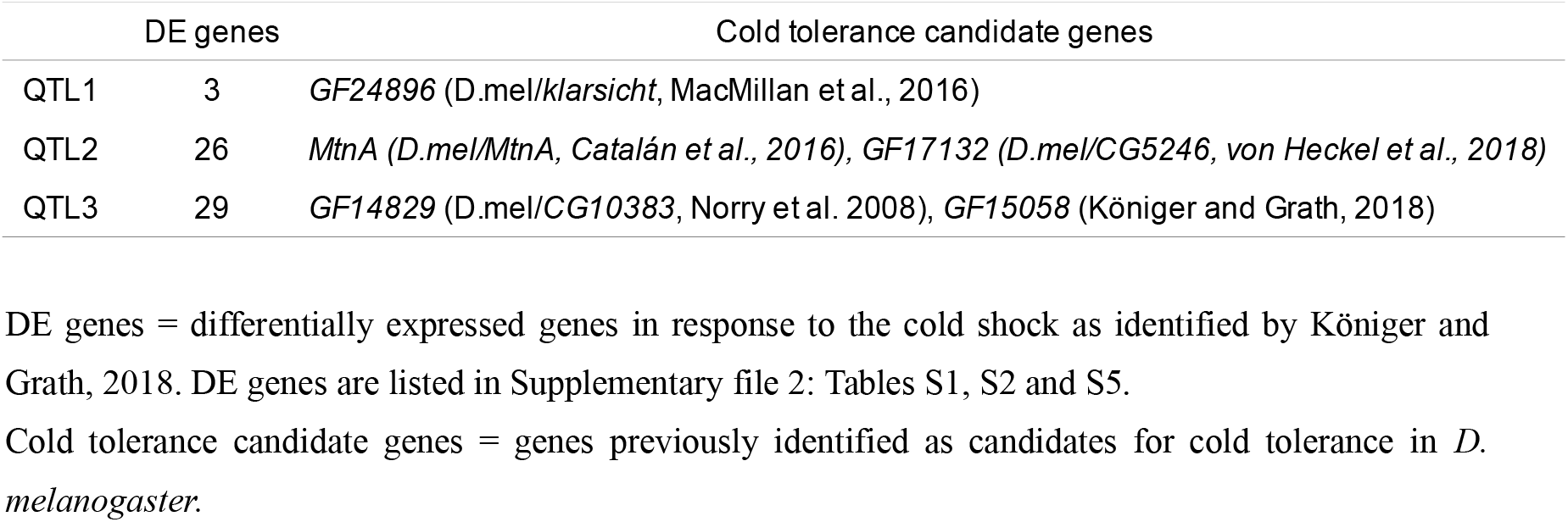
Cold tolerance candidate genes within QTL regions

QTL1 spanned 140 kb and contained eleven protein coding genes (Supplementary file 2: Table S1). There was no enrichment of KEGG pathways or GO terms. However, three of the eleven genes were previously identified to be differentially expressed in response to a cold shock (Supplementary file 2: Table S1, (Königer and Grath, 2018). Two of them, *GF24884* (ortholog of *p130CAS*) and *GF24880* (ortholog of *Phosphoinositide-dependent kinase 1*) were upregulated in both phenotypes after the cold shock and one of them, *GF24896* (ortholog of *klarsicht*) was exclusively upregulated in the cold-tolerant phenotype only. Klarsicht was previously reported as upregulated in cold-acclimated flies of *D. melanogaster* (MacMillan et al., 2016).

QTL2 spanned 1.0 Mb and contained 138 protein coding genes which were enriched in one molecular function, “serine-type endopeptidase activity” (GO:0004252) and one biological process, “intracellular cholesterol transport” (GO:0032367) (Supplementary file 2: Table S3). Out of the 138 genes, 26 were previously identified as differentially expressed in response to a cold shock. Among them, nine genes were upregulated and five genes were downregulated in both phenotypes (see Supplementary file 2: Table S2, and (Königer and Grath, 2018)). In the cold-tolerant phenotype, one gene, *GF17809* (ortholog of *Archease*) was exclusively upregulated and one gene, *GF17856* (ortholog of *Niemann-Pick type C-2c*) was exclusively downregulated. In the cold-sensitive phenotype, one gene, *GF17176* (ortholog of *aluminum tubes*) was exclusively upregulated and nine genes were exclusively downregulated (see Supplementary file 2: Table S2, and (Königer and Grath, 2018).

Nine genes drove the enrichment in the GO category “serine-type endopeptidase activity” (see Supplementary file 2: Table S3). All of them were located in the downstream region of QTL2 at 6,515,565 – 6,527,729 bp and adjacent to one another (Figure 7). Seven of these genes were also differentially expressed in response to cold shock. Among them was *GF17132*, which was upregulated in both phenotypes and its ortholog in *D. melanogaster* showed a significant interaction of phenotype and cold shock (Supplementary file 2: Table S2, von Heckel et al., 2016).

**Figure 7.**
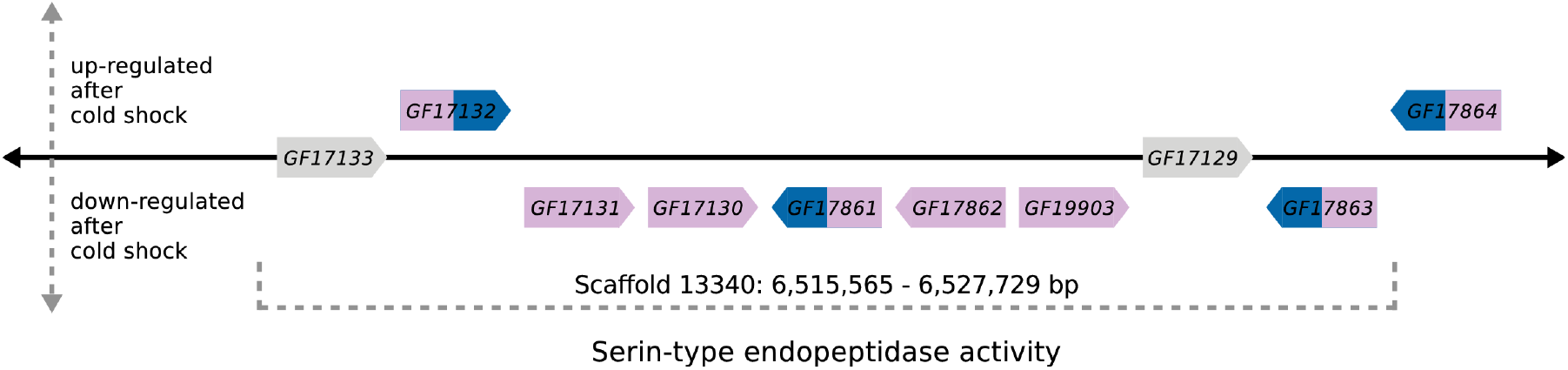
Schematic illustration of a genomic region within QTL2 that contains nine genes of the enriched GO category “serine-type endopeptidase activity” (see also Supplementary file 2: Table S2 and S3). Genes were differentially expressed in response to the cold shock in either the cold-sensitive (slow) phenotype alone (genes shown in pink color) or in both phenotypes, cold-sensitive and cold-tolerant (fast) (genes shown in blue and pink color). Genes that were not differentially expressed are shown in grey. Gene lengths and distances between genes are not drawn to scale.

QTL2 also contained the gene *Metallothionein A* (*MtnA*) which caught our attention because it is involved in metal ion homeostasis and in its *D. melanogaster* ortholog, an InDel polymorphism is associated with local adaptation to oxidative stress upon migration out of Sub-Saharan Africa into Europe (Catalán et al., 2016). *MtnA* was downregulated in response to cold in *D. melanogaster* (von Heckel et al., 2016) but not in *D. ananassae* (Königer and Grath, 2018).

QTL3 spanned 1.2 Mb and contained 110 protein coding genes which were enriched in three molecular functions: “sequence-specific DNA binding” (GO:0043565), “ATPase activity” (GO:0016887) and “phosphotransferase activity, alcohol group as acceptor” (GO:0016773) and one KEGG pathway: “Hippo signaling pathway – fly” (dan04391) (Supplementary file 2: Table S6). Out of the 110 genes, 29 were previously identified as differentially expressed in response to a cold shock (Table 4). Among them, 12 genes were upregulated and seven genes were downregulated in both phenotypic groups, cold-tolerant and cold-sensitive (see Supplementary file 2: Table S5, (Königer and Grath, 2018)). In the cold-tolerant phenotype, two genes, *GF15043* (ortholog of *CG31974*) and *GF14846* (ortholog of *bicoid stability factor*) were exclusively upregulated and two genes, *GF15020* (ortholog of *ABC transporter expressed in trachea*) and *GF14865* (ortholog of *CG11454*) were exclusively downregulated. In the cold-sensitive phenotype, five genes were exclusively downregulated but there were no exclusively upregulated genes. One of the five downregulated genes was *GF15058* (ortholog of *CG10178*), which was one out of two genes with a significant interaction of phenotype and cold shock. The function of *GF15058* is unknown but it is predicted to have UDP-glycosyl-transferase-activity (Marchler-Bauer et al., 2015).

## Discussion

We used a panel of 86 recombinant inbred advanced intercross lines (RIAILs) and 1,400 ddRAD markers to map QTL that underlie natural variation in cold tolerance among two fly strains of *D. ananassae* from a population in Bangkok, Thailand. The recovery time segregated significantly between the two founder strains. CCRT in the cold-sensitive strain was about twice as high as in the cold-tolerant strain. This observation was of particular interest to us because in *D. melanogaster*, previous studies found such differences in the CCRT phenotype only among different populations that inhabit different thermal habitats (David et al., 1998; von Heckel et al., 2016).

The CCRT phenotypes of the mapping population were distributed on a continuum (Figure 3), which was a first important indicator that we were looking at more than one dominant causal allele. Furthermore, three RIAILs recovered faster than the Fast parental strain and 19 of the RIAILs recovered slower than the Slow parental strain, indicating that there was interaction among the parental alleles or loci in the recombinant genotypes of the mapping population.

The three identified QTL for CCRT explain as much as 64% of the variance in the phenotype. This proportion is equal to a previous mapping experiment for CCRT in *D. melanogaster*, in which three QTL explained 64% of the variance for CCRT in an intercontinental set of recombinant inbred lines (Norry et al., 2008). The founder strains for this mapping population were sampled from Denmark and Australia and thus from two geographically different thermal environments. Another study (Morgan and Mackay, 2006) also identified three QTL for CCRT in *D. melanogaster* in a set of recombinant inbred lines derived from two laboratory strains that differed significantly for the phenotype. In this mapping population, the three loci explained 25% of the phenotypic variance for CCRT. While two of the reported QTL for CCRT in *D. melanogaster* co-localized across these two studies, none of the reported candidate genes co-localize with the QTL intervals in *D. ananassae* (this study).

It needs to be noted that, in general, QTL confidence intervals should be considered as support regions rather than absolute boundaries (Broman and Sen, 2009). Further, the causal genetic variants may be located anywhere within these intervals.

Compared to sequencing of pooled samples (Pool-sequencing), RAD-based approaches come at the cost of marker density, especially in crossing designs with low genetic differentiation between the founder strains and low levels of linkage disequilibrium (Futschik and Schlötterer, 2010). Thus, to increase the mapping resolution and to expand the genetic map, we generated a mapping population in which five generations of intercrosses allowed for a sufficient number of crossover events (Pollard, 2012). Subsequently, we used stringent cutoffs for potential sequencing errors and distorted loci. This step certainly increased the robustness of the identified loci, but came at the cost of chromosomal coverage, as many smaller genomic scaffolds were excluded from the analysis at this step. It is therefore possible that our results do not cover all potential QTL. However, the reduction of genome complexity that results from RAD-sequencing has two major benefits. First, it is more cost-effective than whole-genome sequencing of individual samples, allowing for a larger number of samples to be analyzed and consequently for greater statistical power to detect QTL. Second, it is more accurate than whole-genome Pool-sequencing (Catchen et al., 2017; Cutler and Jensen, 2010).

Combining the identified intervals with two previous transcriptome analyses in *D. ananassae* (Königer and Grath, 2018) and *D. melanogaster* (von Heckel et al., 2016) and additional cold tolerance studies in *D. melanogaster* (MacMillan et al., 2016; Norry et al., 2008; Ramnarine et al., 2019) allowed us to narrow down the list of potentially causal genes in *D. ananassae* and to identify common candidate genes in both species. From the combined data, we identified three types of candidates:

I. The expression profile of *GF15058* (*D.mel*/*CG10178*) in QTL3 is directly associated with a difference in the CCRT phenotype in *D. ananassae*. *GF15058* was one out of two genes that responded to the cold shock in a phenotype-specific way (Königer and Grath, 2018). Its function was inferred from electronic annotation to be uridine diphosphate (UDP) glycosyltransferase activity. UDP-glycosyltransferases (UGTs) are membrane-bound enzymes that are located in the endoplasmatic reticulum and catalyze the addition of a glycosyl group from a uridine triphosphate (UTP) sugar to a small hydrophobic molecule. Therefore, UGTs play an essential role in maintaining homeostatic function and detoxification and are known as major members of phase II drug metabolizing enzymes (Bock, 2015). The cold shock led to a downregulation of *GF15058* in the Slow strains but not in the Fast strains. However, the Fast genotype at QTL3 is transgressive, i.e., it increases CCRT. Thus, if *GF15058* was indeed one of the causal factors, our results suggest that keeping transcript abundance at a constant level after the cold shock is so costly for the organism that it slows down recovery.
II. The expression profile of *GF17132* (*D.mel*/*CG5246*) in QTL2 is directly associated with a difference in the CCRT phenotype in *D. melanogaster*, where it showed a significant interaction of phenotype and cold shock (von Heckel et al., 2016). It was also differentially expressed in response to the cold shock in *D. ananassae* (Königer and Grath, 2018). Moreover, *GF17132* belongs to a cluster of genes that code for serine peptidases in QTL2 (Figure 7). Serine peptidases are involved in proteolysis, i.e., they catalyze the hydrolysis of peptide bonds (Attrill et al., 2016; Ross et al., 2003). This process plays a central role in the immune response of insects (De Gregorio et al., 2001) and serine proteases were suggested previously to be involved in the cold stress response as well (Vermeulen et al., 2013).
III. We identified three more genes that have been associated with thermotolerance in experiments other than the transcriptome analyses: *MtnA* (*D.mel*/*MtnA*), *GF24896* (*D.mel/klarsicht)* and *GF14829* (*D.mel/CG10383*). The gene *MtnA* in QTL2 codes for metallothionein A which promotes resistance to oxidative stress. It binds heavy metals and neutralizes reactive oxygen and nitrogen species (Ruttkay-Nedecky et al., 2013). Exposure to cold leads to an increased abundance of free radicals, thereby inducing oxidative stress (Williams et al., 2014). In *D. melanogaster*, a 49 bp deletion in the 3’UTR of *MtnA* is associated with its transcriptional upregulation and with increased tolerance to oxidative stress (Catalán et al., 2016). The frequency of this polymorphism in natural populations follows latitudinal clines, suggesting that upregulation of *MtnA* is favored in temperate environments (Ramnarine et al., 2019). However, a direct link between cold stress and oxidative stress is yet to be established in drosophilids (Plantamp et al., 2016). *MtnA* was downregulated after the cold shock in both phenotypes of *D. melanogaster* (von Heckel et al., 2016) and not differentially expressed in *D. ananassae*. Moreover, a previous sequence analysis of *MtnA* in *D. ananassae* reported the 3’UTR deletion polymorphism as absent in 110 strains that were sampled in tropical and temperate regions around the world (Stephan et al., 1994).

The gene *GF24896* (*D.mel/klarsicht*) in QTL1 is expressed in a wide range of tissues, where it interacts with microtubules and promotes evenly spaced positioning of nuclei. Knock-out of *klarsicht* in muscle cells impairs locomotion and flight (Elhanany-Tamir et al., 2012) – functions that are also disabled during chill coma. The gene was reported previously to be upregulated with cold-acclimation in *D. melanogaster (MacMillan et al., 2016)*. In *D. ananassae*, *GF24896* is upregulated after the cold shock in Fast strains but not in Slow strains (Königer and Grath, 2018), suggesting a potential contribution of this gene to faster recovery from cold exposure.

Lastly, the gene *GF14829* (*D.mel*/*CG10383*) in QTL3 is involved in the regulation of glycosylphosphatidylinositol metabolism. After the cold shock, it is upregulated in Fast and Slow strains of *D. ananassae* and in Slow strains of *D. melanogaster*. Interestingly, over-expression of *CG10383* increases lifespan in *D. melanogaster (Paik et al., 2012).* It was also the only gene within all three QTL for CCRT in *D. ananassae* that mapped to a heat-tolerance QTL in *D. melanogaster* (Norry et al., 2008). In the face of the transgressive nature of QTL3, potential allelic effects resulting in trade-offs between CCRT, heat-resistance and lifespan should be investigated in both species.

In conclusion, we identified three large-effect QTL for recovery from cold exposure in *D. ananassae*. Combining the present results with previous results obtained from *D. melanogaster* allowed us to shed light on commonalities and differences in the genetic basis of cold tolerance between these two species from different phylognetic lineages that have independently expanded their thermal ranges an became successful human commensals. The combined data point at the five above mentioned genes as candidates for recovery from cold exposure. These genes serve as the groundwork for more detailed analyses such as loss-of-function experiments to establish a link between genotype and phenotype in both species.

